# SAMP1 Facilitates Importin-Mediated Nuclear Translocation to Activate the PKA/CREB Pathway and Hepatic Gluconeogenesis

**DOI:** 10.1101/2025.11.17.688768

**Authors:** Marta Columbaro, Elisa Schena

## Abstract

**Background & Aims:** Hepatic gluconeogenesis is pathologically elevated in type 2 diabetes (T2DM). Although the PKA/CREB axis is a central regulator, the mechanisms fine-tuning its activity, particularly nuclear events, remain elusive. This study investigates the role of the inner nuclear membrane protein SAMP1/TMEM201 in this process.

**Methods:** SAMP1 expression was assessed in diabetic (db/db) mice. Gain- and loss-of-function studies were performed in vivo (via AAV8-mediated hepatocyte-specific manipulation in mice) and in vitro (in primary mouse hepatocytes). Mechanisms were probed using co-immunoprecipitation, Western blotting, ELISA, and pharmacological inhibition.

**Results:** Hepatic SAMP1 was upregulated in db/db mice. Overexpression of SAMP1 exacerbated hyperglycemia and glucose intolerance, enhanced gluconeogenic gene expression (Pck1, G6pc), and increased glucose output. Conversely, SAMP1 knockdown attenuated these effects. Mechanistically, SAMP1 interacted with Importinα, facilitating its nuclear translocation. This led to enhanced CREB phosphorylation and activation of gluconeogenic genes, an effect abolished by the CREB inhibitor KG-501.

**Conclusions:** SAMP1 is a novel critical enhancer of hepatic gluconeogenesis. It functions by promoting Importinα-mediated nuclear import of PKA, thereby amplifying the PKA/CREB pathway. Targeting SAMP1 represents a promising strategy for curbing excessive hepatic glucose production in T2DM.

## Introduction

Hepatic gluconeogenesis, the process by which the liver synthesizes glucose, is central to maintaining fasting blood glucose levels. In Type 2 Diabetes Mellitus (T2DM), this pathway becomes hyperactivated, leading to excessive glucose production and contributing significantly to fasting hyperglycemia, a hallmark of the disease^1^. The regulation is complex, involving hormonal signals, key transcription factors, and enzymatic activities.

The canonical pathway involves hormones like glucagon, which, during fasting, activates the PKA/CREB signaling cascade. This leads to the phosphorylation of the transcription factor CREB (cAMP response element-binding protein) and the de-repression of its coactivator CRTC2 (CREB regulated transcription coactivator 2)^2^. This complex then binds to the promoters of key gluconeogenic genes, such as PCK1 (encoding Phosphoenolpyruvate carboxykinase, PEPCK) and G6PC (encoding Glucose-6-phosphatase, G6Pase), driving their expression. Insulin normally counteracts this process, but insulin resistance in T2DM blunts this effect^3^.

SAMP1, also known as Transmembrane Protein 201 (TMEM201), is an integral inner nuclear membrane protein that has garnered significant research interest due to its role in cellular structure maintenance and its implications in cancer and angiogenesis^4–6^. The protein’s location at the nuclear envelope positions it as a potential key regulator of nuclear integrity and cellular signaling pathways, with emerging evidence linking its dysfunction to broader systemic effects. In this article, we discovered that this protein may play a significant role in liver gluconeogenesis and the PKA signaling pathway.

## Results

### 1. SAMP1/TMEM201 Expression is Upregulated in the Livers of Diabetic Models

To investigate the potential role of SAMP1 in metabolic disorders, we first examined its expression in the livers of leptin receptor-deficient (db/db) induced mouse models of type 2 diabetes (Fig.1A). Immunoblotting analysis revealed that SAMP1 protein levels were significantly elevated in this model compared to their respective controls (Fig.1B). This upregulation suggests a potential link between SAMP1 and the dysregulated metabolic state characteristic of diabetes.

**Fig. 1.**
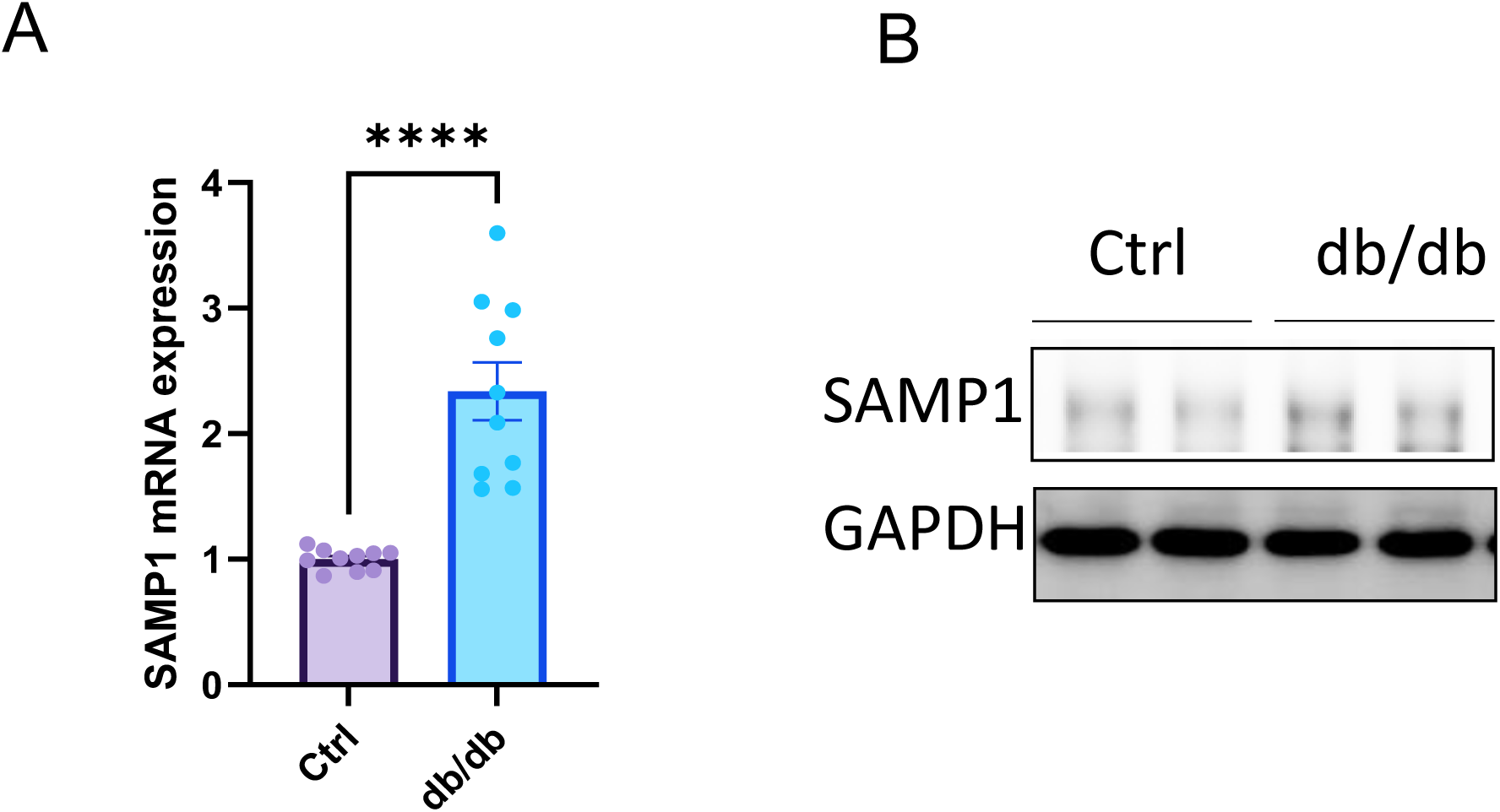
Hepatic SAMP1 is upregulated in vivo. A. SAMP1 mRNA level is increased in livers of db/db mice. B. SAMP1 protein level is increased in livers of db/db mice.

### 2 Overexpression of SAMP1 promotes Hepatic gluconeogenesis in Vitro

To assess the impact of SAMP1 on the regulation of gluconeogenesis in hepatocytes, we modulated SAMP1 expression using Adv-Vector and Adv-SAMP1 in Primary mouse hepatocyte and monitored glucose production changes in response to glucagon. Interestingly, overexpression of SAMP1 could alter glucose production in primary mouse hepatocyte following glucagon administration (Fig. 2A). Consistent with the results, Additionally, RT-PCR analyses confirmed the increased expression of G6PC, and PCK1 in the Adv-SAMP1 group, providing further support for our findings (Fig. 2B-2C). These results confirmed that SAMP1 promoted hepatic gluconeogenesis in vitro under glucagon stimulation

**Fig. 2.**
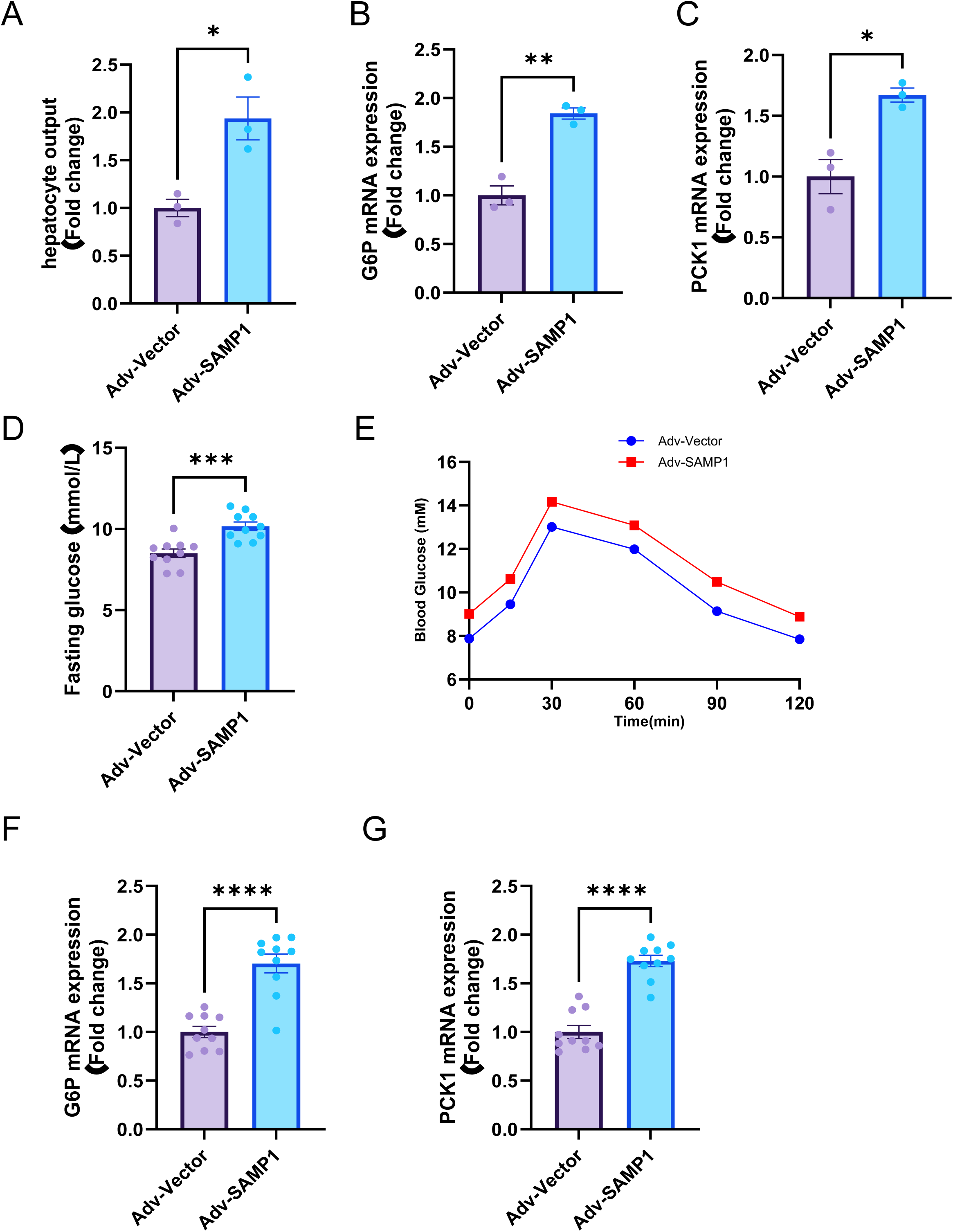
SAMP1 promotes gluconeogenesis in vitro and in vivo. A. SAMP1 overexpression increases glucose production in mouse primary hepatocytes. B. SAMP1 overexpression upregulates G6p mRNA levels in mouse primary hepatocytes. C. SAMP1 overexpression upregulates Pck1 mRNA levels in mouse primary hepatocytes. D. SAMP1 overexpression in mice increases fed blood glucose and impairs PTT. E. SAMP1 overexpression in mice impairs PTT. F. SAMP1 overexpression in mice upregulates G6p mRNA levels. G. SAMP1 overexpression in mice upregulates Pck1 mRNA levels.

### 3 Overexpression of SAMP1 promotes Hepatic gluconeogenesis in Vivo

Hepatocyte-specific SAMP1 overexpression mice were obtained through tail vein injection of AAV8-TBG- SAMP1. The fed blood glucose was increased in the AAV8-TBG-SAMP1 group compared to the AAV8-TBG-SAMP1 group (Fig. 2D). Hepatocyte-specific overexpression of SAMP1 results in impaired GTT in mice (Fig. 2E). Furthermore, the gluconeogenesis-related gene was significantly activated in liver. we observed a significant increase in the expression of the gluconeogenesis genes G6pc (Fig. 2F), and Pck1 (Fig. 2G). Collectively, these findings suggest that SAMP1/TMEM201 may play a role in hepatic gluconeogenesis and have significant implications in systemic glucose metabolism.

### 2. Genetic Knockdown of SAMP1 Attenuates Hepatic Gluconeogenesis

To assess the functional role of SAMP1 in glucose metabolism, we performed genetic knockdown of SAMP1 in primary mouse hepatocytes using adenoviruses expressing shRNA targeting SAMP1. Remarkably, SAMP1 deficiency led to a significant reduction in glucose output under glucagon-stimulated conditions (Fig. 3A). Consistent with this functional change, the mRNA expression of key gluconeogenic enzymes, glucose-6-phosphatase (G6pc) (Fig. 3B) and phosphoenolpyruvate carboxykinase (Pck1) (Fig. 3C), was markedly downregulated in SAMP1-knockdown hepatocytes (Fig. 3D). These results indicate that SAMP1 is required for efficient hepatic gluconeogenesis.

**Fig. 3.**
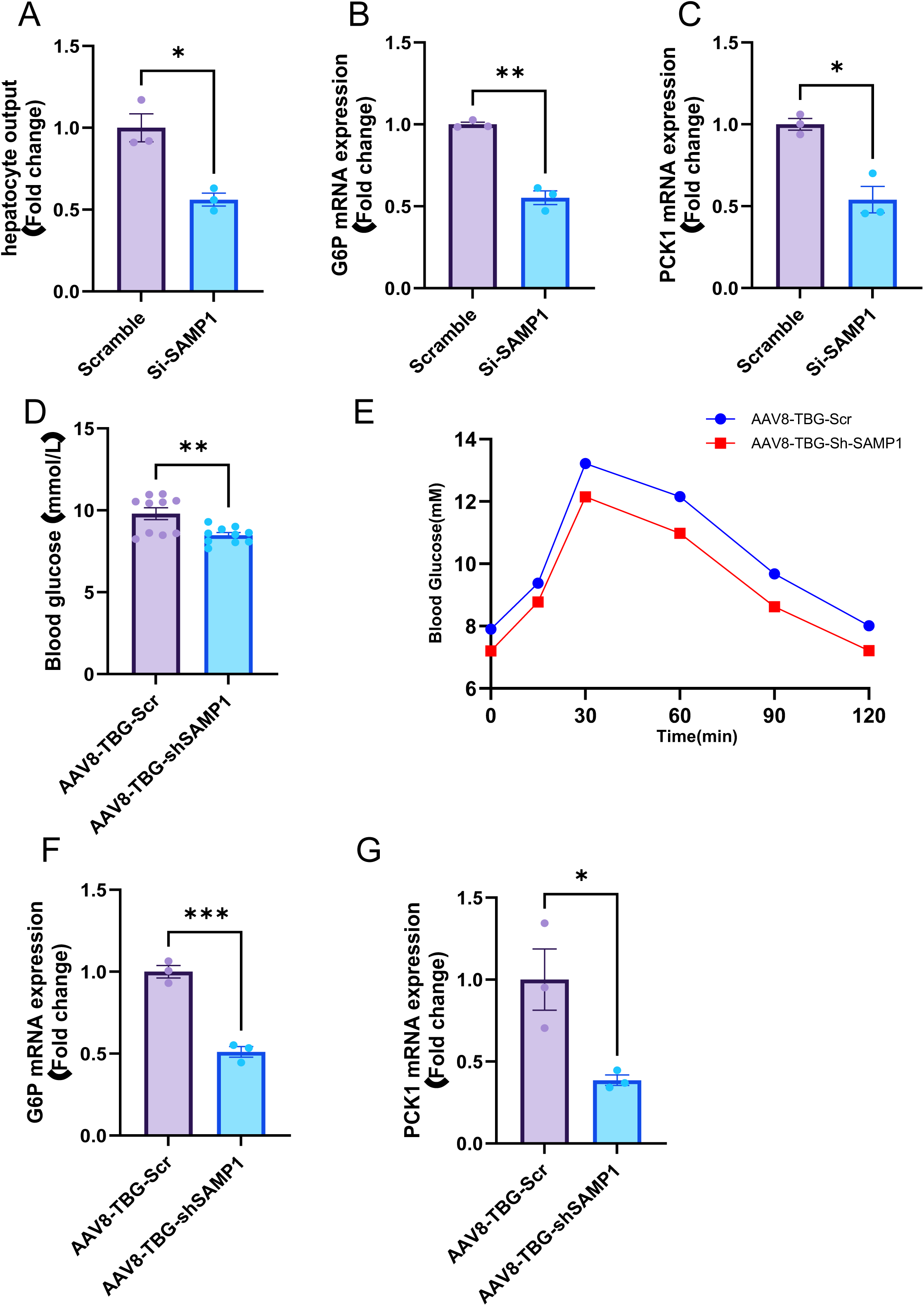
Hepatocyte-specific SAMP1 knock-down inhibited hepatic gluconeogenesis in C57BL/6 mice. A. SAMP1 knock-down decreases glucose production in mouse primary hepatocytes. B. SAMP1 knock-down decreases G6p mRNA levels in mouse primary hepatocytes. C. SAMP1 knock-down decreases Pck1 mRNA levels in mouse primary hepatocytes. D.SAMP1 knock-down in mice decreases fed blood glucose. E. SAMP1 knock-down in mice showed improved PTT. F. SAMP1 knock-down in mice downregulates G6p mRNA levels. G. SAMP1 knock-down in mice downregulates Pck1 mRNA levels.

### 3. Hepatocyte-Specific SAMP1 Knockout Improves Glucose Homeostasis In Vivo

Next, AAV8-TBG-shSAMP1 was employed to interfere with the expression of SAMP1 in the liver of mice to observe its effect on hepatic gluconeogenesis. In the AAV8-TBG-shSAMP1 group, fed blood glucose levels were decreased (Fig. 3D), PTT results were improved (Fig. 3E), and the expression of PCK1 and G6PC in the liver were decreased compared to the AAV8-TBG-SAMP1 group (Fig. 3F,3G). Together, these findings confirmed that downregulating the expression of SAMP1 could reduce hepatic gluconeogenesis

## 4. SAMP1 Interacts with importinα and Regulates the PKA/CREB pathway

Given that SAMP1 is an inner nuclear membrane protein, we hypothesized it might function by regulating the compartmentalization of key metabolic regulators. The Importin family of nuclear transport receptors serves as a fundamental component of the cellular machinery that regulates the movement of molecules between the cytoplasm and the nucleus including PKA^7^. The core mechanism involves the recognition of specific signal sequences on cargo proteins and their facilitated translocation through the Nuclear Pore Complex (NPC)^8^. Co-immunoprecipitation assays in 293T identified importinα as a novel SAMP1-interacting partner (Fig. 4A). Overexpression and SAMP1 knockdown will affect the intranuclear concentration of PKA (Fig. 4B-C). Furthermore, cellular experiments demonstrated that overexpression of SAMP1 promotes CREB phosphorylation (Fig. 4D), and SAMP1 knockdown significantly impaired the glucagon-induced CREB phosphorylation (Fig. 4E). Simultaneously, the CREB phosphorylation inhibitor KG501 could block the regulation by SAMP1 (Fig. 4F).

**Fig. 4.**
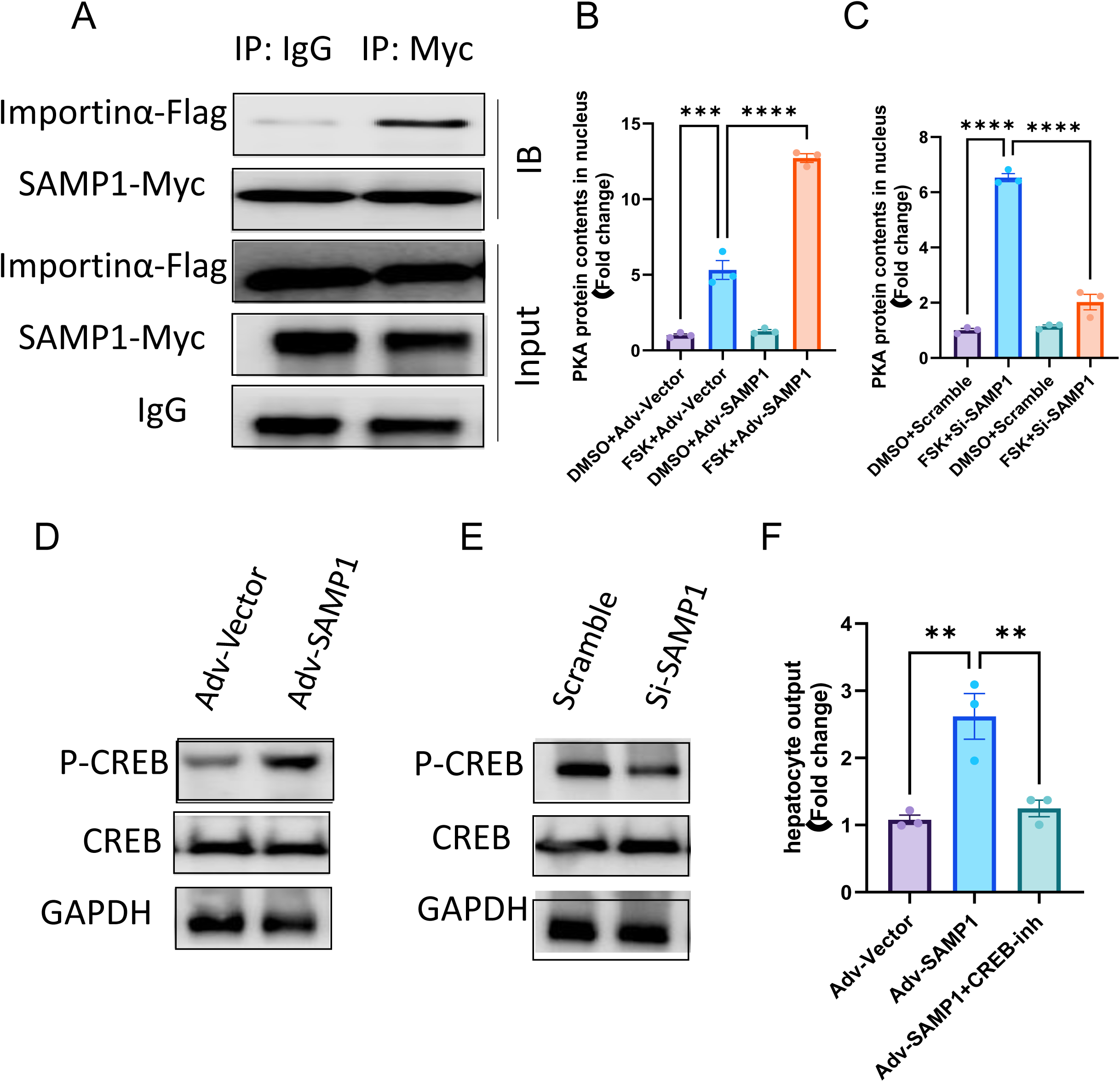
Hepatocyte-specific SAMP1 knock-down inhibited hepatic gluconeogenesis in C57BL/6 mice. A. Representative CO-IP analyses of the interaction between SAMP1 and PKA in 293T cells using exogenous Myc-SAMP1 and Flag -PKA. PKA concentration in cell nucleus of 293T cells PKA concentration in DMSO+Adv-Vector, FSK+Adv-Vector, DMSO+Adv-SAMP1, FSK+Adv-SAMP1 groups. B. PKA concentration in cell nucleus of 293T cells in DMSO+Scramble, FSK+Si-SAMP1, DMSO+Scramble, FSK+Si-SAMP1 groups. C. Western blot analyses p-CREB (S133), CREB and GAPDH in mouse primary hepatocytes from Adv-Vector group and Adv-SAMP1 group. D. Western blot analyses p-CREB (S133), CREB and GAPDH in mouse primary hepatocytes from Scramble and Si-SAMP1 group. E. Glucose production in mouse primary hepatocytes stimulated by glucagon in mouse primary hepatocytes from Adv-Vector group, Adv-SAMP1 group, and Adv-SAMP1+CREB-inh.

## Discussion

In this study, we identify the inner nuclear membrane protein SAMP1/TMEM201 as a novel and critical regulator of hepatic gluconeogenesis. We demonstrate that SAMP1 expression is elevated in the livers of diabetic db/db mice and that its manipulation directly impacts glucose homeostasis: overexpression of SAMP1 exacerbates hyperglycemia and gluconeogenic gene expression, while its knockdown attenuates these processes. Mechanistically, we uncover a previously unrecognized function for SAMP1 in facilitating the nuclear translocation of PKA, thereby potentiating the PKA/CREB signaling cascade. Our findings propose a model wherein SAMP1, by interacting with Importinα, acts as a molecular scaffold that enhances the efficiency of PKA’s nuclear transport, leading to amplified CREB phosphorylation and the transcriptional activation of key gluconeogenic genes like Pck1and G6pc.

Our data introduce SAMP1 as a key player in this spatial regulation. By physically interacting with both PKA and Importinα, SAMP1 appears to facilitate the active nuclear import of PKA, a process that may be more efficient and regulated than passive diffusion. This model is consistent with the growing paradigm that compartmentalized cAMP/PKA signaling is essential for specificity, and SAMP1 represents a crucial component of the nuclear compartmentalization machinery for this pathway. Our discovery that SAMP1 interacts with Importinα provides a direct molecular link between an inner nuclear membrane protein and the canonical nuclear import machinery in the context of metabolic signaling. The Importin family is responsible for the nucleocytoplasmic transport of numerous proteins.

The finding that SAMP1 promotes the nuclear accumulation of PKA suggests that it may act as an adaptor or a facilitator that enhances the recognition of PKA by the Importinα/β complex. This mechanism adds a layer of regulation that ensures a robust and rapid nuclear PKA signal specifically in response to gluconeogenic stimuli, such as glucagon. This is particularly relevant in the context of T2DM, where chronic overactivation of gluconeogenesis is a major driver of fasting hyperglycemia. The upregulation of SAMP1 in diabetic models suggests that this pathway may be hyperactive in the disease state, making SAMP1 a potential therapeutic target. The broader implications of our findings connect with other regulatory mechanisms of the PKA/CREB axis.

In conclusion, we have identified SAMP1/TMEM201 as a novel enhancer of hepatic gluconeogenesis that functions by bridging the Importin-mediated nuclear transport machinery to the PKA/CREB signaling pathway. This study not only expands our understanding of the spatial regulation of cAMP/PKA signaling but also nominates SAMP1 as a potential target for therapeutic intervention in T2DM. Strategies aimed at inhibiting SAMP1 could offer a means to suppress pathological hepatic glucose output without completely abrogating the essential functions of the PKA/CREB pathway in other contexts. Future studies investigating the structural basis of the SAMP1-Importinα-PKA interaction and the metabolic effects of SAMP1 inhibition in adult diabetic animal models will be crucial steps toward validating its therapeutic potential.

## Materials And Methods

Male C57BL/6 mice (8-week-old) and male db/db mice (8-week-old) were acquired from Charles River Laboratories. To generate mice model featuring hepatocyte-specific overexpression and knockdown of the SAMP1, Adeno-associated virus 8 (AAV8)-TBG- SAMP1or AAV8-TBG-sh SAMP1 were administered via tail vein into C57BL/6 or db/db mice, respectively. 4 weeks post AAV8 injection.

Animals were housed under humane conditions with controlled temperature of 22-24° C, relative humidity of 40-60%, a 12-hour light/dark cycle from 7:00 to 19:00, and ad libitum access to food and water. The above-mentioned mice were sacrificed at 12 weeks of age, blood and tissue samples were collected for further experiments. Animal studies were reviewed and approved by the IRCCS Institutial Care and Use Committee, and were in accordance with the Animal Welfare Act and ARRIVE guidelines.

### Glucose and pyruvate tolerance tests

pyruvate tolerance tests (PTT) were conducted in C57BL/6 (4 weeks after viral injection). For the PTT, mice were injected intraperitoneally with sodium pyruvate (2g/kg) after overnight fasting (5:00 P.M. to 9:00 A.M.), and blood glucose levels were measured at 0, 30-, 60-, 90-, and 120-minutes post-injection.

### Cell culture and treatment

Mouse primary hepatocytes (MPHs) were isolated from 6-week-old male C57BL/6 mice as previously reported^9^, and cultured in DME/F12 medium. Cells were incubated at 37°C in a 5% CO2 incubator. For transfection experiments, MPHs were transfected with Adv-*SAMP1* or Adv-sh*SAMP1* for 48 hours and then subjected to subsequent experiments.

### Glucose production assay

Primary hepatocytes were washed three times with PBS and then incubated in glucose and phenol red-free DMEM containing 20 mM sodium lactate and 2 mM sodium pyruvate for 4 hours. Simultaneously, saline, glucagon (100nM, HY-P0082, MedChemExpress), were administered separately to observe the effects of these treatments on glucose production. Glucose levels in the culture medium were measured using a glucose assay Kit (MAK181, Sigma-Aldrich), and the total glucose production was adjusted for protein content.

### RT-PCR

Total RNA was extracted from animal tissues or cells using TRIzol reagent (9109, Takara). Reverse transcription was carried out using the ScriptSure Reverse Transcription Kit (G3326, Takara). Subsequent quantitative PCR reactions were performed using the Takara SYBR Green qPCR Kit (6215A, Takara) and measurements were conducted on a Jena Real-Time PCR System (Analytik Jena). The expression levels of target genes were normalized to those of GAPDH.

### Western blot analysis

Total proteins were isolated from animal tissues or cells using the RIPA lysis buffer (P0013B, Roche), supplemented with a phosphatase inhibitor or protease inhibitor cocktail (4693132001, 4906837001, Roche). Protein concentration was determined using the BCA Protein Assay Kit. Equal protein amounts were separated by 10% SDS-PAGE and transferred to PVDF membranes (IPVH00010, Millipore). Membranes were blocked using a blocking solution, followed by overnight incubation at 4°C with primary antibodies. After three TBST washes, the membranes were incubated with HRP-conjugated secondary antibodies at room temperature for 1 hour. Membrane visualization was done using the Jena Imaging System (Analytik Jena).

### Measurement of Protein Kinase A (PKA) Levels in Cell Lysates by Enzyme-Linked Immunosorbent Assay (ELISA)

Cells were lysed using an ice-cold lysis buffer. To preserve PKA integrity and activity, the buffer was supplemented with protease inhibitors and phosphatase inhibitors (e.g., activated sodium orthovanadate) prior to use. Adherent cells were washed twice with ice-cold phosphate-buffered saline (PBS) and then scraped on ice. Non-adherent cells were pelleted by centrifugation. The cell pellet was resuspended in the supplemented lysis buffer (typically 1 mL per 10C cells) and incubated on ice for 30 minutes with occasional vertexing to ensure complete lysis. The lysate was then centrifuged at 14,000 × g for 15 minutes at 4°C to pellet cellular debris. The resulting supernatant (cell lysate) was carefully collected. For the ELISA, the lysate was typically diluted at least 1:10 in the assay buffer to ensure the measured OD value fell within the linear range of the standard curve. Aliquots of the lysate were stored at -70°C to avoid repeated freeze-thaw cycles, which can degrade protein activity.

Protein Kinase A (PKA) levels in cell lysates were quantified using a commercial mouse PKA sandwich ELISA kit, strictly following the manufacturer’s protocol. The assay is based on a pre-coated plate immobilized with specific anti-PKA capture antibodies. In brief, standards and prepared cell lysate samples were added to the wells, followed by the addition of a horseradish peroxidase (HRP)-conjugated detection antibody. After a series of incubation and washing steps to remove unbound components, tetramethylbenzidine (TMB) substrate was added for color development. The enzymatic reaction was stopped with sulfuric acid, and the absorbance was measured at 450 nm within 15 minutes using a microplate reader. The concentration of PKA in the samples was determined by interpolating the optical density (OD) values from a standard curve generated with known concentrations of standards.

### Cellular Fractionation: Isolation of Cytoplasmic and Nuclear Extracts

The separation of cytoplasmic and nuclear fractions from cultured cells was performed using a modified differential detergent and centrifugation method to study the subcellular localization of proteins of interest. All procedures were carried out on ice or at 4°C to preserve protein integrity and inhibit protease activity. **1. Cell Collection and Washing:** Adherent cells were washed twice with ice-cold phosphate-buffered saline (PBS, pH 7.4) and subsequently scraped into a cold PBS solution. Suspension cells were collected by centrifugation at 300 × g for 5 minutes. The cell pellets were washed twice with PBS to remove residual culture medium.**2. Cytoplasmic Extraction:** The cell pellet was resuspended in a hypotonic lysis buffer (e.g., 10 mM HEPES, pH 7.4, 10 mM KCl, 1.5 mM MgClC, 1 mM DTT) supplemented with 0.1% Nonidet P-40 (NP-40) and a protease/phosphatase inhibitor cocktail immediately before use. The cell suspension was incubated on ice for 10-15 minutes to allow for plasma membrane lysis while keeping the nuclear envelope intact. The lysate was then centrifuged at 3,000 × g for 5 minutes at 4°C. The resulting supernatant, containing the cytoplasmic fraction, was carefully transferred to a new pre-chilled tube. **3. Nuclear Extraction:** The pellet, containing the intact nuclei, was washed once with the hypotonic buffer (without detergent) to remove any residual cytoplasmic contamination. The nuclear proteins were then extracted by resuspending the pellet in a high-salt nuclear extraction buffer (e.g., 20 mM HEPES, pH 7.4, 400 mM NaCl, 1.5 mM MgClC, 0.2 mM EDTA, 10% glycerol, 1 mM DTT) with protease/phosphatase inhibitors. The suspension was vigorously vortexed and incubated on ice for 30-40 minutes with intermittent mixing to facilitate the extraction of nuclear proteins. The insoluble debris was pelleted by centrifugation at 14,000 × g for 15 minutes at 4°C. The clear supernatant, representing the nuclear fraction, was collected.

### Statistical Analysis

All the quantified data are expressed as the means ± SD. An unpaired Student’s t-test was performed to analyze the differences between the two groups. For multiple groups, one-way ANOVA was carried out, followed by the Bonferroni post hoc test. Statistical analyses were performed with GraphPad Prism 9.0 software. A value of p<0.05 was considered significant.

## Contribution

**Marta Columbaro** : Writing- original draft, Data curation, Software, Methodology, **Elisa Schena**: Conceptualization, Funding acquisition, Supervision, Writing – review & editing

## Acknowledgments

The authors thank patients and clinicians for donating samples, the Italian Network for Laminopathies for support and discussion and Aurelio Valmori for the technical assistance.

